# A systematic identification of resistance determinants to antisense antibiotics suggests adaptation strategies dependent on the delivery peptide

**DOI:** 10.1101/2024.10.29.620885

**Authors:** Adam J. Mulkern, Thu-Hien Vu, Linda Popella, Tobias Kerrinnes, Svetlana Ðurica-Mitić, Lars Barquist, Jörg Vogel, Marco Galardini

**Affiliations:** Institute for Molecular Bacteriology, TWINCORE Centre for Experimental and Clinical Infection Research, a joint venture between the Hannover Medical School (MHH) and the Helmholtz Centre for Infection Research (HZI), Hannover, Germany; Cluster of Excellence RESIST (EXC 2155), Hannover Medical School (MHH), Hannover, Germany; Department of Biology, University of Oxford, Oxford, United Kingdom; Institute of Molecular Infection Biology, University of Würzburg, Würzburg, Germany; Cluster for Nucleic Acid Therapeutics Munich (CNATM), Munich, Germany; Helmholtz Institute for RNA-based Infection Research (HIRI), Helmholtz Centre for Infection Research (HZI), Würzburg, Germany; University of Toronto, Mississauga, Ontario, Canada; University of Würzburg, Faculty of Medicine, Würzburg, Germany

## Abstract

The rise of antimicrobial resistance (AMR) among human pathogenic microbes is a serious threat to global health, calling for the development of novel treatment strategies. Antibiotics based on programmable antisense oligomers (asobiotics) offer an attractive solution to the “arms-race”, as their specificity can be quickly updated and tailored to target resistant bacteria. In order to understand the genetic architecture of resistance to asobiotics, we employed laboratory evolution assays to identify mutations that decrease susceptibility to antisense peptide nucleic acid (PNA) against four major gram-negative pathogens: *Escherichia coli, Klebsiella pneumoniae, Salmonella enterica*, and *Pseudomonas aeruginosa*. We observed that the reduction in susceptibility upon asobiotics treatment was dependent on the specific cell penetrating peptide (CPP) being conjugated to the PNAs, suggesting that reduced uptake is a common adaptation strategy only against the (KFF)_3_-K CPP. We in fact observed that *sbmA* was frequently mutated in all tested species when treated with (KFF)_3_-K conjugated PNAs. We further identified mutations related to translation, peptide transport and cell envelope, which provide new hypotheses on cellular response to CPP-PNAs conjugates. Furthermore, for (RXR)_4_XB-*acpP* we observed a modest increase in resistance only when mutations in the PNA binding site were induced, which could easily be bypassed by changing the PNA sequence. These findings indicate that the specific identity of the CPP used plays a key role in determining its robustness against the evolution of resistance, and that laboratory evolution can illuminate the remaining gaps in our knowledge on the mechanisms of action of asobiotics.

## Introduction

Antimicrobial resistance (AMR) poses a significant threat to global public health, necessitating the urgent development of new therapeutic strategies^1^. The introduction of new conventional antibiotics is inevitably followed by the emergence and spread of resistance, which seems to indicate that outpacing the evolution of resistance might not be a realistic strategy^2^. Sequence-based therapeutics, such as short antisense oligomers (ASO) could instead be quickly adapted to circumvent resistance determinants, thanks to their programmable nature, a feature that has been proven to be effective with mRNA vaccines updates^3^. In the context of antimicrobial therapy, asobiotics (*i*.*e*. ASOs that are designed to interfere with bacterial growth) are designed to target the mRNA of essential bacterial genes with sequence complementarity to the translation initiation region. ASOs are thus thought to modulate protein production by sterically blocking ribosome binding and preventing translation initiation^4–7^. Different ASO modalities have been developed^8^, with peptide nucleic acids (PNAs) and peptide phosphorodiamidate morpholino oligomers (PPMOs) being commonly used to target bacteria. PNAs/PPMOs are stabilized and protected from nuclease degradation by the introduction of a neutrally charged pseudo-peptide backbone. To facilitate their delivery into bacterial cells, PNAs/PPMOs are commonly conjugated to cell-penetrating peptides (CPPs). Commonly used CPPs include (KFF)_3_K and (RXR)_4_XB, which are thought to utilize distinct mechanisms for translocation across the bacterial membranes^9^. Previous studies have shown that the (KFF)_3_K-aided delivery of PNAs requires the inner membrane protein *S* bmA for transport ^10–14^, which itself transports the unconjugated PNA after (KFF)_3_K has been degraded by a yet unidentified periplasmic protease. On the other hand, (RXR)_4_XB has been shown to operate through a membrane-potential dependent mechanism^15^.

Even though there have been substantial improvements in the development of asobiotics, for instance by defining the optimal ASO sequence characteristics^16^, a broad understanding of the mechanisms of resistance induced by treatment with asobiotics is currently lacking. Based on what is known about their mode of action, it can be expected that resistance mechanisms will fall in roughly three categories: reduced binding to the target site, impaired uptake, and other mechanisms. Mutations in the ASO binding region have indeed been proven to affect asobiotics efficacy, even though they have not yet been reported to spontaneously emerge upon treatment^17^. Spontaneous mutations affecting active uptake have instead been previously described, specifically in the antimicrobial peptides transporter protein SbmA which have been shown to confer resistance to (KFF)_3_K-mediated delivery of PNAs and PPMOs into bacterial cells^10,11^. If these resistance mutations were frequent across clinically relevant species and asobiotic formulations, it might be possible to focus on designing broad-spectrum treatments that induce these mutations with low frequency^18^; the empirical data is however currently lacking. Moreover, a systematic view of asobiotics resistance mutations would help to generate new hypotheses on alternative uptake pathways and the cellular response to the stress induced by the interference with the translation of essential genes. This deeper understanding would in turn also allow for the design of formulations that are more resilient to the emergence of resistance.

In this study, we aimed to systematically investigate the resistance to CPP-PNA formulations using *in vitro* experimental evolution. We tested four CPP-PNA formulations across four clinically relevant gram-negative bacterial pathogens. We found that the choice of CPP has a large influence on the magnitude of induced resistance, and we determined which genetic variants are induced in response to treatment, some of which seem to be related to the cellular adaptation to the asobiotics mode of action. These results suggest that in order to develop programmable sequence-based antimicrobials that are resilient to the evolution of resistance, great consideration must be taken in selecting the most appropriate delivery mechanism.

## Results

### (KFF)_3_K induces the highest increase in resistance in a “drug selection ramp”

To induce resistance we implemented a “drug selection ramp”, an *in vitro* laboratory evolution method that has been previously described^19,20^. This serial passage assay applies a gradual increase in selection pressure, starting from a sub-inhibitory concentration (half of the minimum inhibitory concentration, MIC) and doubling the concentration every four passages, reaching four times the MIC in the last passage (**Figure 1**). We applied this assay on six individual strains belonging to four clinically relevant pathogen species: *E. coli*, *K. pneumoniae*, *S. enterica*, and *P. aeruginosa*. For *E. coli* we tested three distinct strains: the MG1655 reference, the KEIO *sbmA* knockout strain (with the BW25113 background, mutant ID JW0368), and the uropathogenic (UPEC) strain 536^21^. The Δ*sbmA* mutant was chosen for the role of the encoded inner membrane transporter in the uptake of (KFF)_3_K-conjugated PNAs into *E. coli*; knocking out this gene renders *E. coli* less susceptible to (KFF) _3_K-PNA treatment^11^. The Δ*sbmA* mutant could therefore provide insights into additional genes influencing PNA activity that are otherwise overlooked due to the predominant role of mutations in *sbmA*.

**Figure 1.**
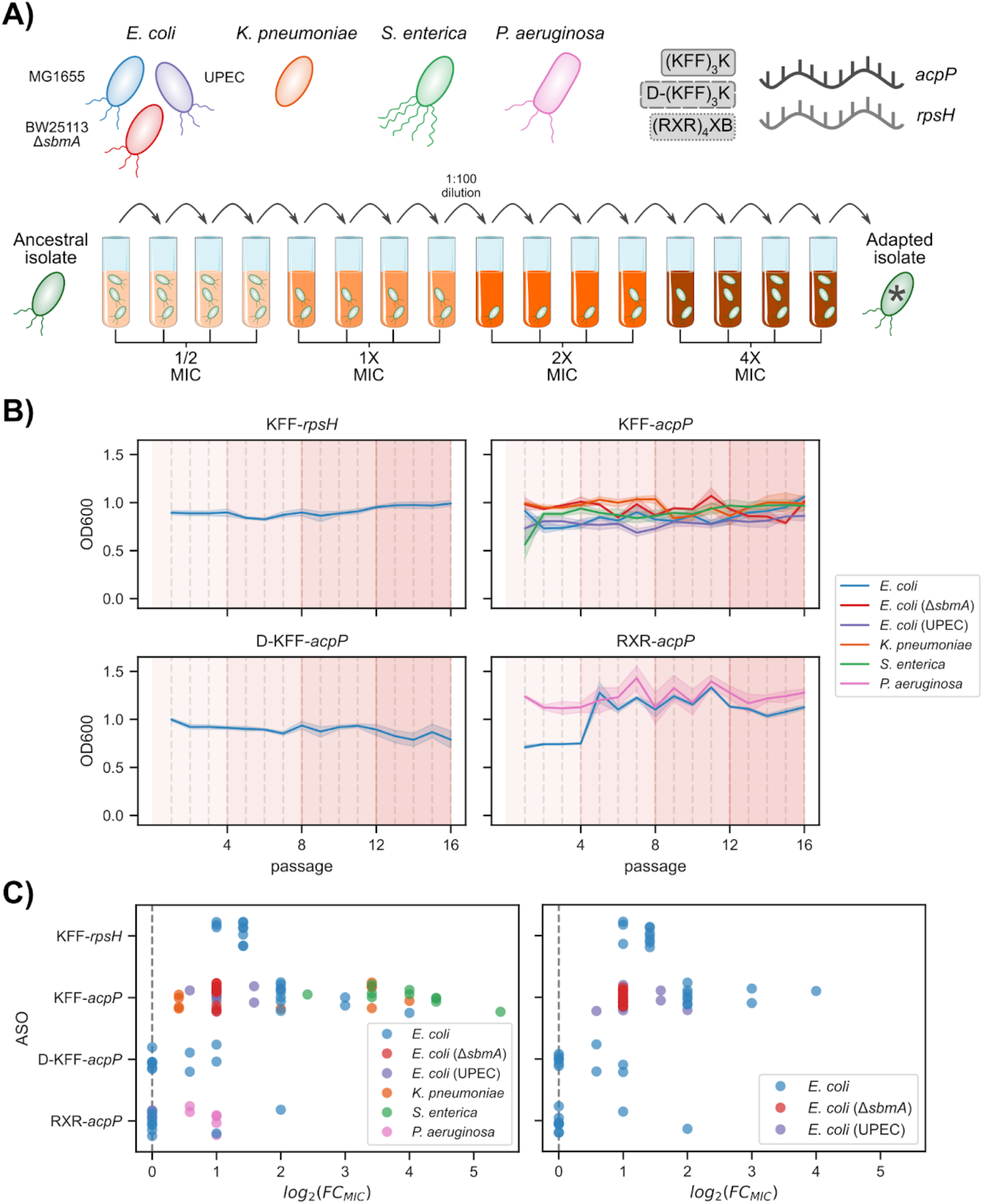
A drug selection ramp to induce resistance to asobiotics. **A**) Schematic representation of the species and asobiotic formulations tested, as well as the serial passaging protocol. **B**) Daily optical density (OD) measurements from the 9 drug selection ramps; the solid line indicates the average OD over the 10 biological replicates, the shaded area the 95% confidence interval. **C**) Fold-change increase in minimum inhibitory concentration (MIC) after the completion of the selection ramp with respect to the ancestral isolate. Each dot indicates the fold-change for a single biological replicate. The right sub-panel shows the measurements only for the three *E. coli* strains.

We tested four different ASO formulations, combining two distinct PNAs, targeting the *rpsH* and *acpP* essential genes, respectively, with three CPPs; we used (KFF) _3_K in its original L-form, its D-form (D-(KFF)_3_K), and (RXR)_4_XB, termed as KFF, D-KFF, and RXR hereafter (**Figure 1A**). We tested KFF-*acpP* on all strains except *P. aeruginosa*, for which we tested a cognate RXR-*acpP* construct; *P. aeruginosa* lacks an *sbmA* homolog, very likely accounting for the ineffectiveness of KFF for PNA delivery for this species^22^. In contrast, all four ASO formulations were tested against the *E. coli* MG1655 reference strain to directly compare them: KFF-*acpP*, D-KFF-*acpP*, KFF-*rpsH*, and RXR-*acpP*. Ten biological replicates were performed for each asobiotic/strain combination, in order to better evaluate the variance in the induced resistance phenotype.

We observed consistent levels of growth during each of the 16-daily 1:100 passages of the drug selection ramp, with limited variability across replicates (**Supplementary Figure 1**), suggesting that each isolate may have developed resistance to the asobiotic formulation by the end of the ramp (**Figure 1B**). We confirmed that on average all drug selection ramps induced resistance by performing susceptibility assays, which revealed increased MIC values between the ancestral and adapted populations. We observed the highest increase in MIC for *S. enterica* adapted to KFF-*acpP*, with an average fold increase of 17.6. On the other hand, we observed the lowest increase for *E. coli* MG1655 adapted to D-KFF-*acpP*, with an average fold increase of 1.3. Strikingly, for *E. coli* MG1655 adapted to RXR-*acpP* we did not record any increase in MIC with respect to the ancestral isolate for 8 out of the 10 adapted populations (**Figure 1C, Supplementary Figure 2**, and**Supplementary Table 1**). When focusing on *E. coli* MG1655, which was tested on all four asobiotic formulations, we observed a clear difference in MIC increase based on the CPPs used, with the two KFF formulations inducing a higher level of resistance (mean MIC fold change of 2.5 and 6 for KFF-*rpsH* and KFF-*acpP*, respectively) when compared to both D-KFF-*acpP* and RXR-*acpP* (1.3 and 1.4 average increase in MIC, respectively). These results prove (KFF)_3_K to be a less resilient choice of CPP for asobiotic delivery which is in line with previous reports showing its low stability and its dependency on *sbmA*-mediated active uptake^13^. We further confirmed this observation by noting that the MIC increase of the Δ*sbmA* mutant was on average higher than that we observed for both D-KFF-*acpP* and RXR-*acpP* (2 versus 1.3 and 1.4, respectively). Therefore, this mutant has additional capacity to increase its resistance on top of the already existing *sbmA* gene deletion.

### (KFF)_3_K asobiotics induce genetic variants in *sbmA* at high frequency across species

To uncover the genetic basis of the observed resistance, we carried out whole-genome sequencing of both the ancestral and adapted experimental populations (**Supplementary Table 2**). On average, we counted 11.4 genetic variants for each biological replicate, with *P. aeruginosa* having the highest number of variants (31.8 for each replicate on average). We recorded the “large deletion” as the most common category of genetic variant across all samples (average of 5.6 deletions per sample). We used the frequency by which each gene was found to be mutated across the 10 biological replicates as a way to identify the most likely determinants of resistance for each asobiotic construct (**Figure 2**, **Supplementary Figure 3**). For this analysis we excluded synonymous single-nucleotide polymorphisms (SNPs), variants in intergenic regions, and those in known pseudogenes.

**Figure 2.**
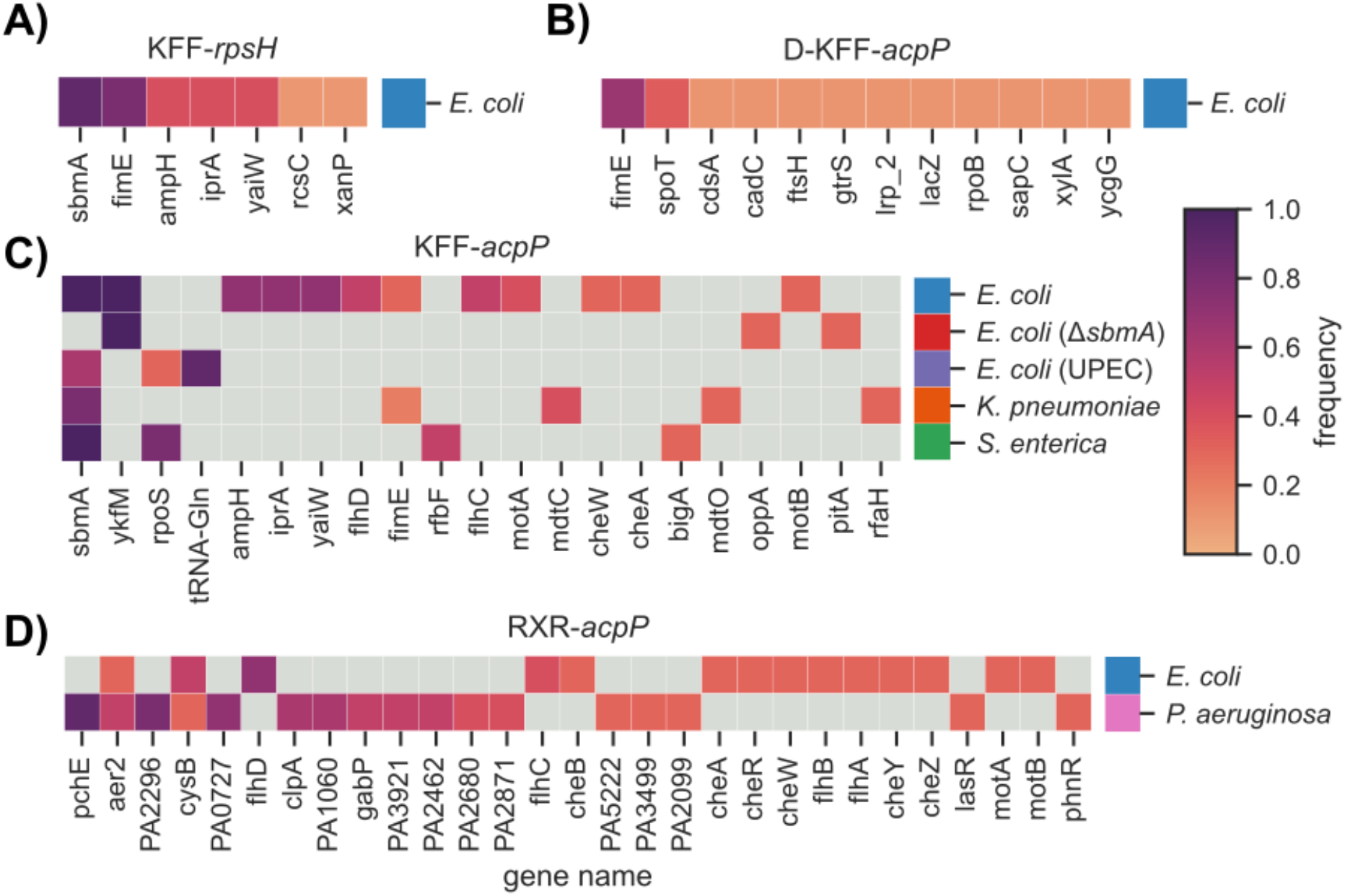
Common genetic variants induced during the drug selection ramps. The heatmaps indicate the proportion of replicates for each strain/asobiotic combination for which we observed genetic variants in each gene. Gray boxes indicate that no variant was observed in the corresponding gene. For KFF-*acpP* and RXR-*acpP*, which each have a list of genes with at least one sample showing a mutation, only genes with a variant in at least three replicates are shown. Genes are sorted by their average frequency across all species, with high to low frequency sorted from left to right.

For all the species adapted to KFF-*acpP*, we observed a high (at least 6 replicates) frequency of likely impactful genetic variants in the *sbmA* gene, which is essential for the transport of KFF-conjugated PNAs into the cell. We observed a similarly high frequency (9 replicates) of variants in the *sbmA* gene for *E. coli* MG1655 adapted to KFF-*rpsH*, suggesting that resistance to KFF in this strain outweighs that to the PNA itself. This result extends previous reports on the role of mutations in *sbmA* in resistance to KFF-PNAs beyond a single strain of *E. coli* and including *S. enterica* and *K. pneumoniae* ^11^. For *E. coli* MG1655 we observed a high frequency of variants in *yaiW* when adapted to both KFF-*acpP* (7 replicates) and KFF-*rpsH* (4 replicates); this gene is co-transcribed with *sbmA*, encodes for an outer membrane protein, and its deletion mutant has been shown to affect the sensibility to antimicrobial peptides, which overall may indicate a functional relationship with the better characterized SbmA^23^. Additionally, variants in the *ykfM* gene were consistently identified across all replicates of both *E. coli* MG1655 and Δ*sbmA* adapted to KFF-*acpP*. This uncharacterized transmembrane gene has been linked to alterations in response to chemical and physical stresses^24^. We observed that *fimE*, involved in the fimbrial biosynthetic process, was frequently mutated in *E. coli* MG1655 adapted to all KFF-conjugated PNAs. FimE has been associated with cell adhesion, and mutations near the *fimA* start codon could influence bacterial attachment and biofilm formation in response to antibiotic exposure^25^.

It has been previously shown how SbmA transports PNAs from periplasmic space into the cytoplasm after KFF has been proteolytically degraded, and how its D-form (D-KFF), by virtue of its resistance to proteolysis, is not dependent on this transport mechanism^11^. Consistent with this model, we did not observe any mutation in *sbmA* when we treated *E. coli* with D-KFF-*acpP* (**Supplementary Figure 10**). We instead observed that D-KFF-*acpP* induced variants in the *spoT* gene in *E. coli* (3 replicates), possibly linking the SpoT-dependent stress response to fatty acid metabolism, which is directly targeted by the *acpP* PNA ^26^. We further identified variants induced in *xanP* in *E. coli* MG1655 treated with KFF-*acpP* (1 replicate,**Supplementary Figure 5**), KFF-*rpsH* (1 replicate,**Supplementary Figure 8**), and RXR-*acpP* (1 replicate, **Supplementary Figure 12**), as well as in the Δ*sbmA* mutant (2 replicates, **Supplementary Figure 9**); this gene encodes for a nucleobase transporter^25^. Additionally, KFF-*acpP* induced variants in the *oppA* gene in *E. coli* Δ*sbmA* (3 replicates), which is involved in peptide transport^27^. In *S. enterica*, the most common mutations were found in *rpoS* (8 replicates), which encodes the RNA polymerase sigma factor, and in *rfbF* (5 replicates), a gene involved in the biosynthesis of the LPS O-antigen in the outer membrane, which has been shown to inhibit PNA efficacy in *E. coli* ^28^. In *K. pneumoniae*, we observed mutations in *mdtC* in 4 replicates. This gene is part of the *mdtABC/tolC* multidrug efflux pump and confers resistance to several antimicrobials^29^. Even though chemical inhibitors of the pump activity have shown no impact on PNA efficacy^28^, the *bamB/tolC* double mutant was found to be more sensitive to KFF-*acpP* in *E. coli* ^30^. We also observed frequent mutations in *fimE* in *K. pneumoniae* when adapted to KFF-*acpP*, similar to what we observed in *E. coli*. Finally, both D-KFF-*acpP* and RXR-*acpP* formulations did not induce a large increase in MIC in most samples (**Figure 1**). Therefore, we hypothesize that the high-frequency mutations observed may not be directly related to resistance to these ASO constructs and might represent “passenger” mutations instead.

### Private mutations are associated with variability in the induced resistance to asobiotic treatment

Given the variability in the MIC fold changes across the different biological replicates, we next focused on the differences between individual samples adapted to the same asobiotic formulation. We looked in particular at *E. coli* MG1655, which was adapted to all four asobiotics, to study the joint variability in phenotypic and genotypic adaptation (**Figure 3A**, and **Supplementary Figures 5-13**). For KFF-*rpsH* we observed a relatively uniform increase in MIC across all replicates (2.5 fold-change over the ancestral on average), and with all isolates encoding a variant in *sbmA*, as well as *fimE* in eight out of ten replicates. Populations of *E. coli* MG1655 adapted to KFF-*acpP* had both higher average MIC increases (6 fold-change over the ancestral) and larger variance, with three replicates increasing their MIC with respect to the ancestral isolate more than eight times. Since all samples developed similar *sbmA* genetic variants, we focused on variants exclusive to three samples with higher MIC increase (**Supplementary Figure 5**). For sample M03, which showed an 8-fold increase in MIC we observed a private mobile genetic element insertion in the *lrhA* gene, a probable transcriptional regulator. For sample M10, which also showed an 8-fold increase in MIC, we observed two private non-synonymous SNPs in the *metK* (V231F) and *ysaA* (W3R) genes, both of which have no known involvement in resistance to asobiotics or the KFF CPP. Lastly, for sample M02 we observed a 16-fold increase in MIC, the above-mentioned variant in *xanP*, and a non-synonymous SNP, not found in any other sample, in codon 246 (H246L) of the *prfB* gene. *prfB* encodes for the peptide chain release factor RF2, which among other functions interacts with ArfA to recover stalled ribosomes^31,32^; mutations at position 246 in particular have been shown to increase termination efficiency^33,34^. Even though the mechanism of action of asobiotics is thought to be the prevention of ribosome binding through steric interference, we found the function of this gene in general, and of mutations at this position in particular, compelling, also in light of the absence of other unique variants. Interestingly, in UPEC adapted to KFF-*acpP*, we have observed a variant in the related gene *prfA* (**Supplementary Figure 7**), specifically a C127F substitution, which further highlights the potential relevance of *prf*-related genes in adaptation to asobiotics treatments.

**Figure 3.**
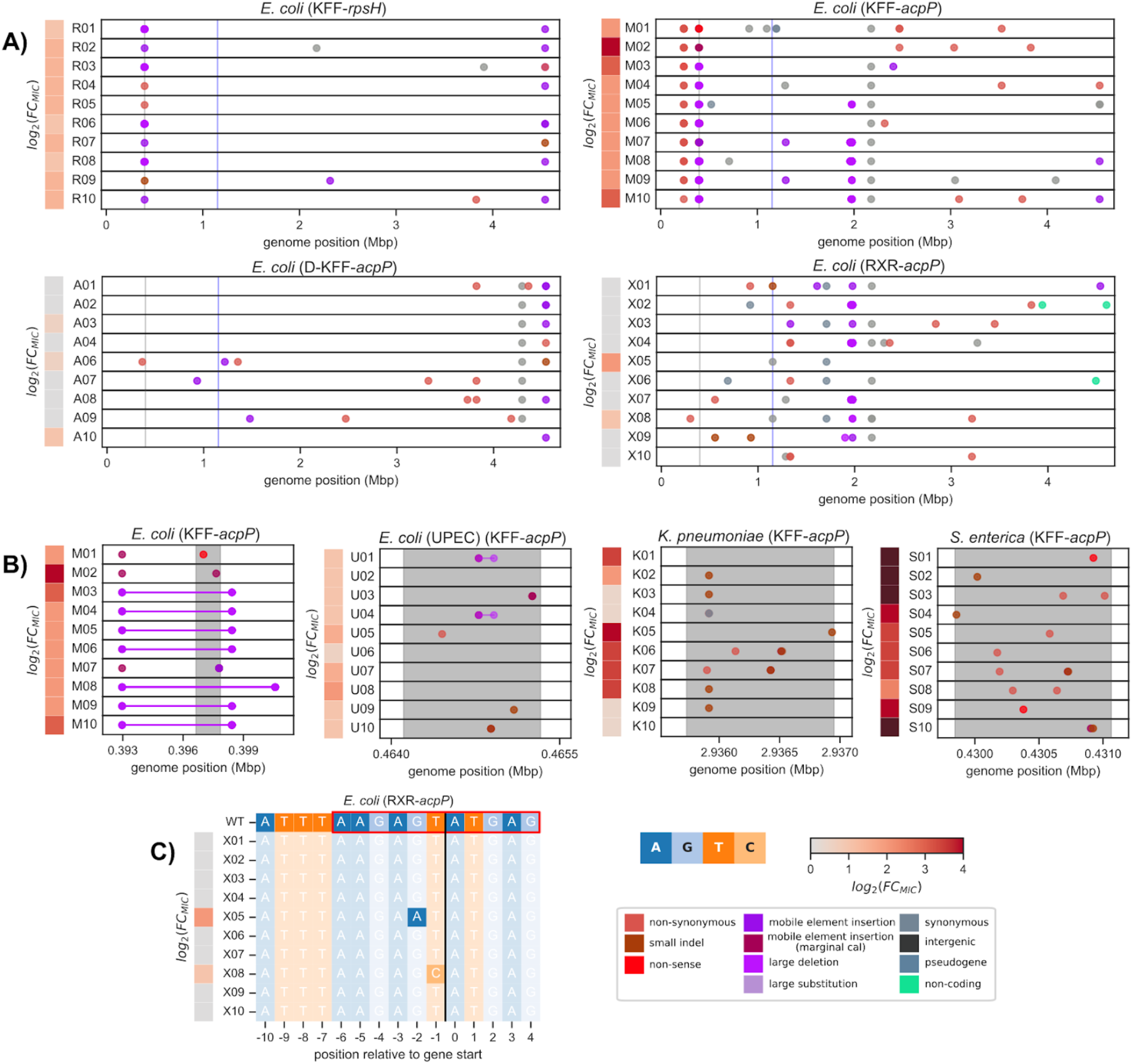
A detailed view on the genome-wide distribution of genetic variants induced by the drug selection ramps. **A**) Genome-wide distribution of genetic variants in each biological replicate for *E. coli* MG1655 after exposure to each of the four asobiotic formulations; the MIC fold-change with respect to the ancestral isolate is shown in gray-to-red heatmaps on the left. The gray and blue vertical lines indicate the location of the *sbmA* and PNA target genes (*acpP* or *rpsH*), respectively. The same visualization for the other five species/ASO formulations can be found in **Supplementary Figure 4. B**) Genetic variants in the *sbmA* gene induced by treating *E. coli* MG1655, UPEC 536, *K. pneumoniae* or *S. enterica* with KFF-*acpP*. The gray shaded area in each panel indicates the location of *sbmA*’s coding sequence. Each subpanel has a different genomic window size in order to accommodate the variants in and around *sbmA*.**C**) Rare genetic variants in *acpP* induced within the RXR-*acpP* targeting window in *E. coli* MG1655; the red box indicates the location of the PNA binding site. The two replicates with SNPs within the target sequence are highlighted, together with their MIC fold-change increase. The vertical black line indicates the start of *acpP*’s coding sequence.

Even though all three tested species adapted to KFF-*acpP* developed mutations in *sbmA*, we observed some variability in the types of induced variants across replicates (**Figure 3B**); for *E. coli* MG1655 we observed large (average 5771 bp) deletions mediated by the IS3 insertion sequence in 7 replicates, 5 of which had the same length (5465 bp) and start and end position on the chromosome. We observed the exact same IS3-mediated deletions in 4 replicates of *E. coli* MG1655 adapted to KFF-*rpsH* (**Supplementary Figure 8**). The deletion variants induced by KFF-*acpP* in UPEC 536 were in contrast rarer (2 replicates) and shorter (both 134 bp). At the other end of the spectrum, we noted that variants induced in both *K. pneumoniae* and *S. enterica* were all short variants such as non-synonymous SNPs (3 replicates for *K. pneumoniae*, 5 for *S. enterica*), non-sense SNPs (2 replicates for *S. enterica*), and small insertion/deletions (InDels, 7 replicates for *K. pneumoniae*, 4 for *S. enterica*).

### A relatively rare mutation within the PNA target window is associated with reduced sensitivity to RXR-*acpP* in *E. coli*

Since RXR-aided PNA uptake is independent from any active transporter machinery, there might be no direct way of developing resistance against the RXR CPP, which likely explains observing less frequent mutations across replicates and the average low increase in MIC. As reported above and in **Figure 1**, we observed that only two *E. coli* MG1655 populations developed a measurable increase in MIC when compared to the ancestral isolate when adapted to the RXR-*acpP* formulation. The most notable genetic variants (**Supplementary Figure 12**) differentiating these samples from the remaining eight was the presence of two SNPs in the center of the RXR-*acpP* binding site, i.e. at position -1 (T>C) and -2 (G>A) for sample X08 (2 fold MIC increase) and X05 (4 fold MIC increase), respectively (**Figure 3C**). While up to two mismatches on the 5’ or 3’ binding target of PNA and PPMO-based asobiotics do not completely abolish its binding capacity, mismatches in the central binding region substantially impair PNA binding^16,17,35^. In both samples we observed a relatively low frequency (∼30%) of these SNPs, which may explain the relatively small increase in MIC. From this result we can conclude that *E. coli* (strain MG1655 in particular) seems to be unable to successfully escape the detrimental effects of RXR- or (D)-KFF-mediated delivery of the *acpP* PNA.

## Discussion

A crucial step in developing new antimicrobial compounds is identifying the resistance mechanisms that will inevitably emerge once these treatments are being used at scale^36^. This step is not only important to forecast the evolution of resistance, but also to elucidate further the mechanism of action of the new compounds. The selective pressure exerted by the antimicrobial is likely to elicit mutations in genes that are directly related to the action of the drug and the cellular response to it. In the case of asobiotics this is of particular importance, as empirical data on their uptake, precise mode of action, and impact on cellular processes is still scarce. Additionally, data on whether clinically relevant species respond in the same way to CPP-PNA conjugates is also currently lacking.

In this study, we have for the first time used a systematic approach to identify the genetic determinants for asobiotics resistance across CPP-PNA formulations and clinically relevant bacterial gram-negative species (*E. coli*, *S. enterica*, *K. pneumoniae*, and *P. aeruginosa*). An important observation from our study is the high frequency of mutations in the *sbmA* gene when using KFF-conjugated PNAs; while these mutations had been previously reported for *E. coli* ^11^, here we have shown how this gene is frequently mutated in two other clinically relevant species such as *S. enterica* and *K. pneumoniae*. Beyond *sbmA*, in *E. coli* treated with KFF-*acpP* we observed frequent variants in *yaiW*, which is co-transcribed with *sbmA* and is localized in the outer membrane; its role in resistance to antimicrobial peptides might give it a role in the uptake of KFF-conjugated PNAs^23^. The downstream gene product YaiW, is in fact a palmitate-modified surface-exposed outer membrane lipoprotein^23^, indicating a possible initial entry mechanism for the KFF-PNAs into the periplasmic space, then followed by a SbmA-dependent transport into the cytoplasm. This potential complementary role of YaiW in KFF-PNAs transport is however yet unproven. Other frequent mutations of note include the *ykfM* gene, encoding an uncharacterised transmembrane protein, suggest an additional possible asobiotic transport mechanism and therefore a potential role in resistance. However, since knockout mutants of this gene show changes in their chemical and physical stress response pathways^24^, this mutation could instead be influencing PNA efficacy indirectly. Furthermore, we observed other species-specific mutations induced by treatment with KFF-*acpP*, including the glutamine-tRNA gene in UPEC 536, *rfbF* in *S. enterica*, and *mdtC* in *K. pneumoniae*. Mutations that might alter translation rates and efficiency, which is a plausible consequence of a mutation in a tRNA gene, and more so in the peptide release factors *prfA* and *prfB*, might ameliorate the reduced rate of translation of the target gene. Both *prfA* and *prfB* are known to influence translation by rescuing stalled ribosomes^31,32,37^; we can therefore hypothesize that mutations in these two genes can confer resistance to PNAs via two mechanisms. The first one could be of an indirect nature, via an overall increased rate of translation, which would also affect the target essential gene, thus counteracting the effect of the PNA’s steric interference. The second one would be via a direct interaction with the ribosome subunits that fail to form a functional complex on the target mRNA because of the PNA binding. The current available information is however insufficient to clarify which of these hypotheses are correct, if any. Regarding mutations in *rfbF*, we hypothesize that they could result in significant alterations to the outer membrane structure, thus potentially impacting the ability for CPPs to bind and penetrate the cell membrane^28^, and in turn, inhibit PNA delivery. Again, these altered membrane properties could have further downstream impacts on the susceptibility of the pathogen not only to asobiotics, but also to other antimicrobials. Similarly, we have observed frequent mutations in *mdtC*,which is part of the *mdtABC* / *tolC* multidrug efflux pump; as *tolC* / *bamB* double knockout mutants in *E. coli* have been shown to have higher susceptibility to KFF-*acpP*, we can anticipate that investigating efflux pumps more closely might further elucidate KFF-PNAs uptake^30^.

It has been previously demonstrated that introducing mutations in the central asobiotic binding site results in its reduced efficacy^16,17^. To the best of our knowledge, it has not been previously reported to happen spontaneously upon treatment. We observed two instances of mutations in the central portion of the asobiotic *acpP* binding site when treating *E. coli* MG1655 with RXR-*acpP*, which coincidentally were the only samples treated with this formulation to develop any measurable increase in their MIC. This striking correlation, and existing *in-vitro* data on this very genomic region^16^, leads us to hypothesize with relatively high confidence that these mutations are the sole cause of the increase in resistance for these two samples. We also use the observation that mutations in the asobiotic binding site were induced in only two samples over the total of 90 evolved lines to hypothesize that mutations in these highly conserved promoter regions might incur a fitness cost; this in turn makes them less likely to emerge as opposed to the other mutations we have observed. Even if the frequency of these mutations was eventually found to be higher in more realistic treatment scenarios, they could be circumvented with relative ease through an update of the PNA sequence, thus making asobiotic a treatment that is resilient to the evolution of resistance.

As we had expected based on theoretical considerations, we have observed that asobiotic treatment induces mutations affecting both uptake and the target sequence itself. For some of the remaining variants we could make hypotheses on their role in resistance (*e.g*.outer membrane composition and efflux), but perhaps more intriguingly on the actual mode of action of this new class of antimicrobials (*i.e*.rescue of stalled ribosomes). Future work should be directed at further characterizing these variants to validate these hypotheses and clarify the molecular mechanisms behind asobiotics action and cellular response, with a special focus on their evolutionary resilience.

## Materials and Methods

### Bacterial strains

The following strains were used in this study: *Escherichia coli* MG1655 (JVS-5709; NCBI GenBank: U00096.3), *Pseudomonas aeruginosa* PAO1 (JVS-11761; NCBI RefSeq: NC_002516.2), *Klebsiella pneumoniae* ICC8001 (kindly provided by Joshua Liang Chao Wong and Gad Frankel, Imperial College London; JVS-13403; NCBI GenBank: CP009208.1), uropathogenic *E. coli* 536 (UPEC 536; JVS-12054; NCBI GenBank: CP000247.1). All strains were streaked on Luria-Bertani (LB) agar plates to obtain single colonies and cultured in non-cation adjusted Mueller-Hinton Broth (MHB, BD Difco™, Thermo Fisher Scientific), at 37°C and 220 rpm shaking.

### Peptide nucleic acids (PNAs)

Peptide-conjugated PNAs (PPNA), directly linked to either (KFF)_3_K, D-(KFF)_3_K or (RXR)_4_XB, were purchased from Peps4LS GmbH (Heidelberg, Germany). The constructs’ quality and purity were verified by mass spectrometry and HPLC, ensuring a purity level above 98%. PPNAs were dissolved in nuclease-free water and heated at 55°C for 5 minutes before use. Concentrations were measured using a NanoDrop spectrophotometer (A260 nm; ThermoFisher, U.S.) and calculated based on the extinction coefficient^38^. The PPNAs were stored at –20°C and reheated at 55°C for 5 minutes before preparing the working dilutions. Throughout, low retention pipette tips and low binding Eppendorf tubes (Sarstedt, Germany) were used.

### Determination of Minimal Inhibitory Concentration (MIC) and Minimal Bactericidal Concentration (MBC)

The MIC values were determined using a modified broth microdilution method^39^, following the guidelines of the Clinical and Laboratory Standards Institute. An overnight bacterial culture was diluted 1:100 in fresh Mueller-Hinton Broth (MHB) and cultivated to an optical density (OD600) of 0.5. This culture was further diluted in MHB to achieve a final concentration of approximately 10^5^ cfu/ml. The diluted bacterial solution was then combined with 10x PPNA working solutions in a 96-well plate (Thermo Fisher Scientific, U.S.), adjusting a final volume of 100 µl per well. Growth was monitored using a BioTek Epoch2NS Microplate Spectrophotometer (Agilent, U.S.), measuring OD600 every 20 minutes for 24 hours at 37°C with continuous double-orbital shaking (237 cpm). The MIC was defined as the lowest PPNA concentration that inhibited visible growth (OD600 < 0.1).

To determine the MBC, immediately following the MIC assessment, the entire contents of wells containing ½x, 1x and 2x MIC were plated onto LB agar. These plates were incubated at 37°C for 18-20 hours, and bacterial growth was recorded to identify the lowest PPNA concentration preventing regrowth. To detect potential slow-growing colonies, the plates were further incubated at room temperature for an additional 72 hours.

### Experimental Evolution

Resistance was induced by subjecting bacterial strains to progressively increasing concentrations of PPNA over 16 daily passages. Initial susceptibility values determined by MIC were used to design a modified ‘evolutionary ramp’ protocol^19,20^, with PPNA concentrations increased every four passages from ½x MIC to 4x MIC. The passages were conducted in 96-well microplates (Greiner Bio-One, Austria) with 10 replicates for each strain and PPNA combination (150 µl total well volume). Plates were incubated for 22 hours at 37°C, and overnight cultures were diluted 1:100 into fresh LB broth medium. Optical density (OD600nm) was measured at the end of each passage. All samples, including both control and treated strain groups, were stored in 25% glycerol at -80°C.

### DNA isolation, Sequencing and Data Analysis

To assess genetic variations in microbial populations from experimental evolution, DNA was extracted from the bulk liquid culture of each 10 replicated treatment groups and one control ancestral group for every combination of PPNA and strain at the final collection point (passage 16). The DNeasy Blood & Tissue Kit (Qiagen, Germany) was used for genomic DNA extraction, following the manufacturer’s protocol for gram-negative bacteria. DNA concentration and purity were measured using a NanoDrop spectrophotometer (dsDNA, A_260_/A_280_ ratio). Library preparation using standard Illumina protocols, and sequencing using the Illumina HiSeq 4000/PE150 platform was performed at Novogene (Cambridge, UK). Variant populations were detected using breseq (v.0.38.2)^40^, a pipeline tool developed for detecting mutations in laboratory evolving microorganisms. The analysis was performed using the polymorphism mode to detect variants present in a non-clonal bacterial population. Mutations were filtered based on their frequency, with the threshold set at 15%. Ancestral isolates, for which we isolated the DNA from a single colony, were instead analyzed using the consensus mode of breseq. For sample M02 we called variants again using snippy (v4.6.0)^41^, as breseq did not annotate the non-synonymous variant in *prfB* correctly due to the presence of a known ribosome slippage site in this gene. For each adapted sample we adopted the following data processing steps on the GenomeDiff files: we first subtracted variants observed in the corresponding ancestral isolate. We then identified the variants present across all 10 replicates for each strain/ASO combination and removed them from each sample; we reasoned that the likelihood of observing the exact same genetic variant over 10 biological replicates would be extremely low, and that those variants would be most likely be present in the ancestral isolate. In order to match genetic variants across strains and species and identify frequently mutated genes we annotated the reference genomes using eggnog-mapper (v2.1.12)^42^, and used the orthologous group identifier to match ortholog genes.

## Code and data availability

Raw sequencing reads have been deposited at the European Nucleotide Archive (ENA), with accession number PRJEB81806. All the scripts used to process the OD readings, MIC values, the outputs of breseq, and generate all figures presented in the manuscript are available as a git repository: https://github.com/microbial-pangenomes-lab/2024_aso_evolution. The version of the code used to generate all the results presented here has been deposited in Zenodo^43^. The code is based on python scripts and Jupyter notebooks, using the following libraries: numpy (v2.0.0)^44^, scipy (v1.14.0)^45^, pandas (v2.2.2)^46^, statsmodels (v0.14.2), matplotlib (v3.9.1)^47^, seaborn (v0.13.2)^48^, and jupyter-lab (v4.2.4)^49^.

## Author contributions

MG, JV, AJM and LP conceived the study, AJM and THV performed susceptibility testing, the laboratory evolution assays and prepared samples for whole genome sequencing, AJM and MG analyzed the data and generated figures, with help from LB in interpreting results from genome sequencing. LP, TK, and SDM sourced the asobiotics and provided guidance on handling them. AJM and MG wrote the manuscript draft, and all authors provided edits and comments.

## Acknowledgements

AJM and MG were supported by the Deutsche Forschungsgemeinschaft (DFG, German Research Foundation) under Germany’s Excellence Strategy - EXC 2155 - project number 390874280. THV was supported by the Hannover Biomedical Research School (HBRS), and the Center for Infection Biology (ZIB), as well as the Graduate School Scholarship Program (GSSP) from the German Academic Exchange Service (DAAD). This work was further supported by funds to JV from a DFG Gottfried Wilhelm Leibniz Award (DFG Vo875-18) and the Bavarian consortium bayresq.net project Rbiotics (JV, LB). JV and LP were supported by the BMBF in the framework of the Cluster4Future program (Cluster for Nucleic Acid Therapeutics Munich, CNATM Project ID: 03ZU1201CA).

## References

1. Murray, C. J. et al. Global burden of bacterial antimicrobial resistance in 2019: a systematic analysis. The Lancet399, 629–655 (2022).

2. Ventola, C. L. The Antibiotic Resistance Crisis. Pharm. Ther. 40, 277–283 (2015).

3. Verbeke, R., Lentacker, I., De Smedt, S. C. & Dewitte, H. The dawn of mRNA vaccines: The COVID-19 case. J. Control. Release Off. J. Control. Release Soc. 333, 511–520 (2021).

4. Good, L. & Nielsen, P. E. Inhibition of translation and bacterial growth by peptide nucleic acid targeted to ribosomal RNA. Proc. Natl. Acad. Sci. 95, 2073–2076 (1998).

5. Good, L. & Nielsen, P. E. Antisense inhibition of gene expression in bacteria by PNA targeted to mRNA. Nat. Biotechnol. 16, 355–358 (1998).

6. Dryselius, R., Aswasti, S. K., Rajarao, G. K., Nielsen, P. E. & Good, L. The translation start codon region is sensitive to antisense PNA inhibition in Escherichia coli. Oligonucleotides13, 427–433 (2003).

7. Popella, L. et al. Comprehensive analysis of PNA-based antisense antibiotics targeting various essential genes in uropathogenic Escherichia coli. Nucleic Acids Res. 50, 6435–6452 (2022).

8. Ghosh, C. et al. A comparative analysis of peptide-delivered antisense antibiotics using diverse nucleotide mimics. RNA N. Y. N30, 624–643 (2024).

9. Inoue, G., Toyohara, D., Mori, T. & Muraoka, T. Critical Side Chain Effects of Cell-Penetrating Peptides for Transporting Oligo Peptide Nucleic Acids in Bacteria. ACS Appl. Bio Mater. 4, 3462–3468 (2021).

10. Puckett, S. E. et al. Bacterial Resistance to Antisense Peptide Phosphorodiamidate Morpholino Oligomers. Antimicrob. Agents Chemother. 56, 6147–6153 (2012).

11. Ghosal, A., Vitali, A., Stach, J. E. M. & Nielsen, P. E. Role of SbmA in the Uptake of Peptide Nucleic Acid (PNA)-Peptide Conjugates in E. coli. ACS Chem. Biol. 8, 360–367 (2013).

12. Howard, J. J. et al. Inhibition of Pseudomonas aeruginosa by Peptide-Conjugated Phosphorodiamidate Morpholino Oligomers. Antimicrob. Agents Chemother. 61, e01938–16 (2017).

13. Yavari, N., Goltermann, L. & Nielsen, P. E. Uptake, Stability, and Activity of Antisense Anti-acpP PNA-Peptide Conjugates in Escherichia coli and the Role of SbmA. ACS Chem. Biol. 16, 471–479 (2021).

14. Ghilarov, D. et al. Molecular mechanism of SbmA, a promiscuous transporter exploited by antimicrobial peptides. Sci. Adv. 7, eabj5363 (2021).

15. Frimodt-Møller, J. et al. Activating the Cpx response induces tolerance to antisense PNA delivered by an arginine-rich peptide in Escherichia coli. Mol. Ther. - Nucleic Acids25, 444–454 (2021).

16. Jung, J. et al. Design and off-target prediction for antisense oligomers targeting bacterial mRNAs with the MASON web server. RNA N. Y. N29, 570–583 (2023).

17. Nanayakkara, A. K., Moustafa, D. A., Pifer, R., Goldberg, J. B. & Greenberg, D. E. Sequence specificity defines the effectiveness of PPMOs targeting Pseudomonas aeruginosa. Antimicrob. Agents Chemother. 67, e0024523 (2023).

18. Wojciechowska, M., Równicki, M., Mieczkowski, A., Miszkiewicz, J. & Trylska, J. Antibacterial Peptide Nucleic Acids—Facts and Perspectives. Molecules25, 559 (2020).

19. Gifford, D. R. et al. Identifying and exploiting genes that potentiate the evolution of antibiotic resistance. Nat. Ecol. Evol. 2, 1033–1039 (2018).

20. San Millan, A., Escudero, J. A., Gifford, D. R., Mazel, D. & MacLean, R. C. Multicopy plasmids potentiate the evolution of antibiotic resistance in bacteria. Nat. Ecol. Evol. 1, 1–8 (2016).

21. Hochhut, B. et al. Role of pathogenicity island-associated integrases in the genome plasticity of uropathogenic Escherichia coli strain 536. Mol. Microbiol. 61, 584–595 (2006).

22. Ghosal, A. & Nielsen, P. E. Potent antibacterial antisense peptide-peptide nucleic acid conjugates against Pseudomonas aeruginosa. Nucleic Acid Ther. 22, 323–334 (2012).

23. Arnold, M. F. et al. Enteric YaiW is a surface-exposed outer membrane lipoprotein that affects sensitivity to an antimicrobial peptide. J. Bacteriol. 196, (2014).

24. Han, X. et al. Escherichia coli genes that reduce the lethal effects of stress. BMC Microbiol. 10, 35 (2010).

25. Tirumalai, M. R. et al. Evaluation of Acquired Antibiotic Resistance in Escherichia coli Exposed to Long-Term Low-Shear Modeled Microgravity and Background Antibiotic Exposure. mBio10, 10.1128/mbio.02637-18 (2019).

26. Battesti, A. & Bouveret, E. Acyl carrier protein/SpoT interaction, the switch linking SpoT-dependent stress response to fatty acid metabolism. Mol. Microbiol. 62, 1048–1063 (2006).

27. Kashiwagi, K., Yamaguchi, Y., Sakai, Y., Kobayashi, H. & Igarashi, K. Identification of the polyamine-induced protein as a periplasmic oligopeptide binding protein. J. Biol. Chem. 265, 8387–8391 (1990).

28. Good, L., Sandberg, R., Larsson, O., Nielsen, P. E. & Wahlestedt, C. Antisense PNA effects in Escherichia coli are limited by the outer-membrane LPS layer. Microbiology146, 2665–2670 (2000).

29. Hernando-Amado, S. et al. Multidrug efflux pumps as main players in intrinsic and acquired resistance to antimicrobials. Drug Resist. Updat. Rev. Comment. Antimicrob. Anticancer Chemother. 28, 13–27 (2016).

30. Yamamoto, K. et al. Scaffold size-dependent effect on the enhanced uptake of antibiotics and other compounds by Escherichia coli and Pseudomonas aeruginosa. Sci. Rep. 12, 5609 (2022).

31. Chadani, Y., Ito, K., Kutsukake, K. & Abo, T. ArfA recruits release factor 2 to rescue stalled ribosomes by peptidyl-tRNA hydrolysis in Escherichia coli. Mol. Microbiol. 86, 37–50 (2012).

32. Shimizu, Y. ArfA recruits RF2 into stalled ribosomes. J. Mol. Biol. 423, 624–631 (2012).

33. Mora, L., Heurgué-Hamard, V., de Zamaroczy, M., Kervestin, S. & Buckingham, R. H. Methylation of bacterial release factors RF1 and RF2 is required for normal translation termination in vivo. J. Biol. Chem. 282, 35638–35645 (2007).

34. Dinçbas-Renqvist, V. et al. A post-translational modification in the GGQ motif of RF2 from Escherichia coli stimulates termination of translation. EMBO J. 19, 6900–6907 (2000).

35. Goltermann, L., Yavari, N., Zhang, M., Ghosal, A. & Nielsen, P. E. PNA Length Restriction of Antibacterial Activity of Peptide-PNA Conjugates in Escherichia coli Through Effects of the Inner Membrane. Front. Microbiol. 10, 1032 (2019).

36. Spagnolo, F., Trujillo, M. & Dennehy, J. J. Why Do Antibiotics Exist? mBio 12, e0196621 (2021).

37. Vivanco-Domínguez, S. et al. Protein synthesis factors (RF1, RF2, RF3, RRF, and tmRNA) and peptidyl-tRNA hydrolase rescue stalled ribosomes at sense codons. J. Mol. Biol. 417, 425–439 (2012).

38. Popella, L. et al. Global RNA profiles show target selectivity and physiological effects of peptide-delivered antisense antibiotics. Nucleic Acids Res. 49, 4705–4724 (2021).

39. Goltermann, L. & Nielsen, P. E. PNA Antisense Targeting in Bacteria: Determination of Antibacterial Activity (MIC) of PNA-Peptide Conjugates. Methods Mol. Biol. Clifton NJ 2105, 231–239 (2020).

40. Deatherage, D. E. & Barrick, J. E. Identification of mutations in laboratory evolved microbes from next-generation sequencing data using breseq. Methods Mol. Biol. Clifton NJ 1151, 165–188 (2014).

41. Seemann, T. snippy: fast bacterial variant calling from NGS reads https://github.com/tseemann/snippy. (2015).

42. Cp, C., A, H.-P., I, L., P, B. & J, H.-C. eggNOG-mapper v2: Functional Annotation, Orthology Assignments, and Domain Prediction at the Metagenomic Scale. Mol. Biol. Evol. 38, (2021).

43. Galardini M. microbial-pangenomes-lab/2024_aso_evolution: Initial submission. Manuscript version 2024: https://zenodo.org/records/14002597.

44. Harris, C. R. et al. Array programming with NumPy. Nature 585, 357–362 (2020).

45. Virtanen, P. et al. SciPy 1.0: fundamental algorithms for scientific computing in Python. Nat. Methods 17, 261–272 (2020).

46. McKinney, W. Data Structures for Statistical Computing in Python - SciPy Proceedings. https://proceedings.scipy.org/articles/Majora-92bf1922-00a (2010).

47. Hunter, J. Matplotlib: A 2D Graphics Environment. https://ieeexplore.ieee.org/document/4160265 (2007).

48. Waskom, M. L. seaborn: statistical data visualization. J. Open Source Softw. 6, 3021 (2021).

49. Perez, F. & Granger, B. IPython: A System for Interactive Scientific Computing | IEEE Journals & Magazine | IEEE Xplore. https://ieeexplore.ieee.org/document/4160251 (2007).

