## Supplementary material for "A systematic identification of resistance determinants to antisense antibiotics suggests adaptation strategies dependent on the delivery peptide"

#### Supplementary tables

- **Supplementary Table 1.** Minimum inhibitory concentration (MIC) values for ASO formulations for each ancestral strain and adapted isolates.
- **Supplementary Table 2.** List of genetic variants identified in each sample adapted to the ASO formulations, after filtering.

### Supplementary figures

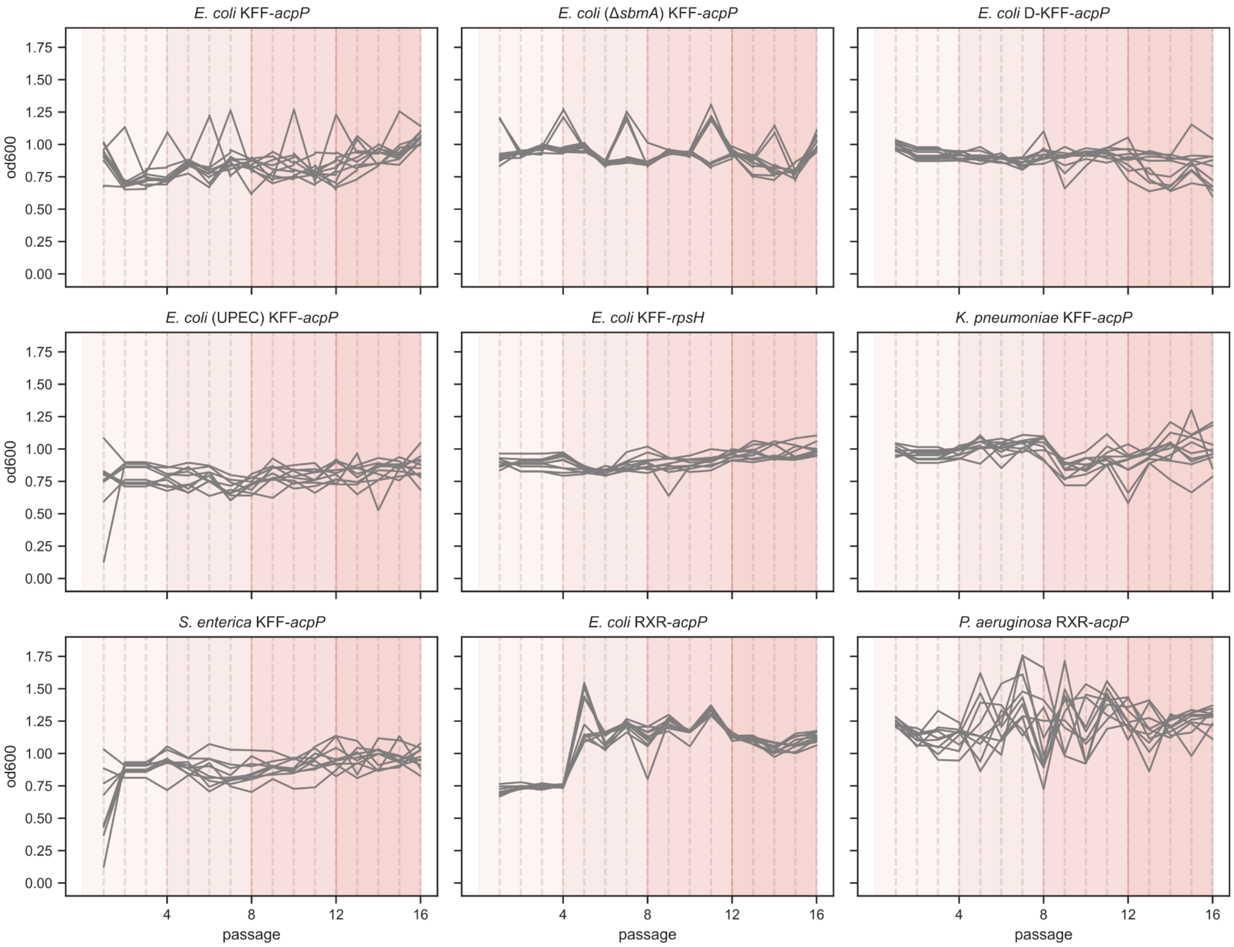


**Supplementary Figure 1. Daily optical density (OD) readings from the 9 drug selection ramps.** Each line indicates the OD readings of one of the 10 biological replicates.

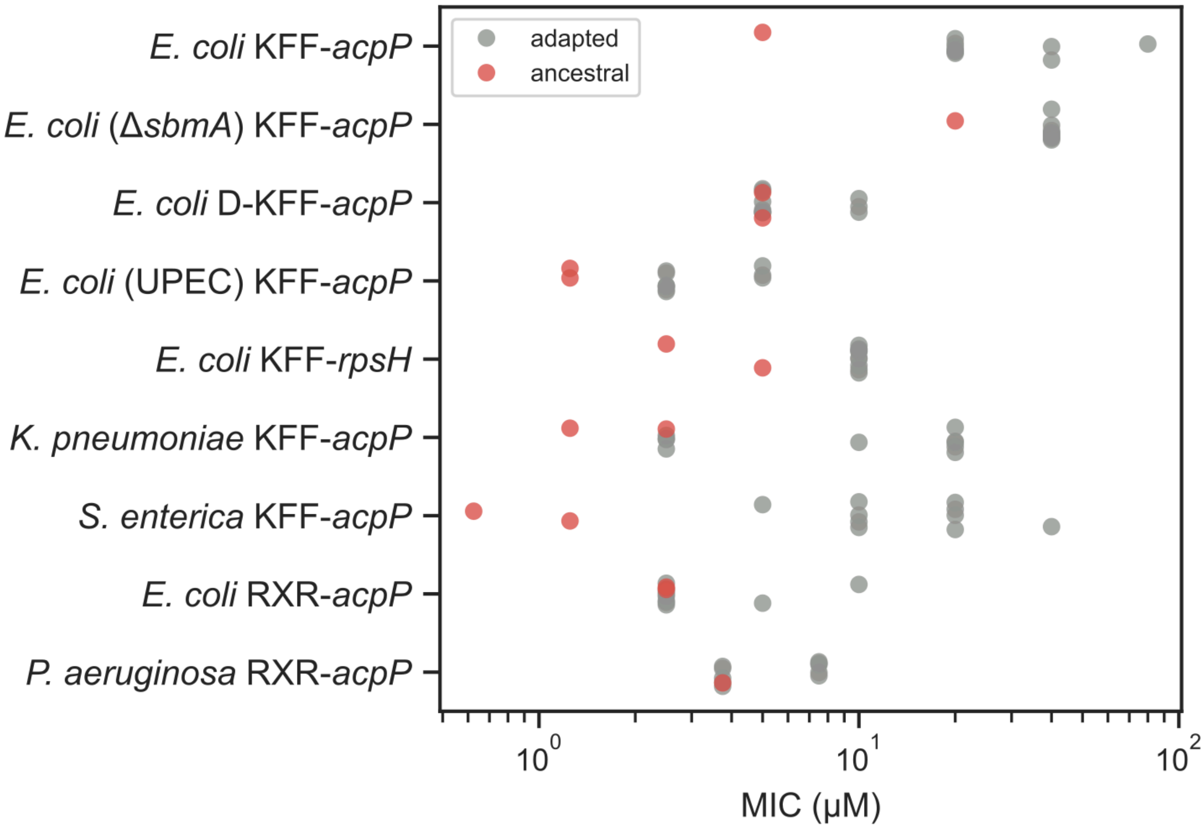


**Supplementary Figure 2. Absolute MIC values for all ancestral and adapted isolates.**

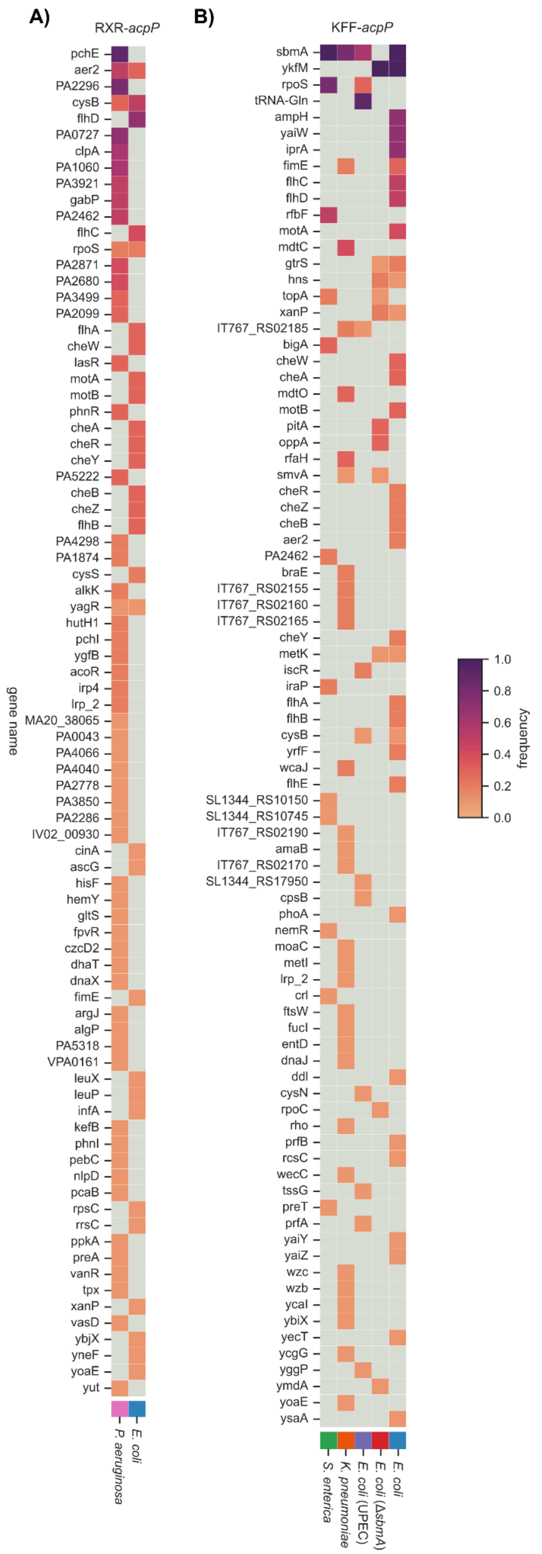


**Supplementary Figure 3. Common genetic variants induced during the drug selection ramps, full list.** For RXR-*acpP* (A) and KFF-*acpP* (B), all genes with at least one genetic variant in one sample are reported.

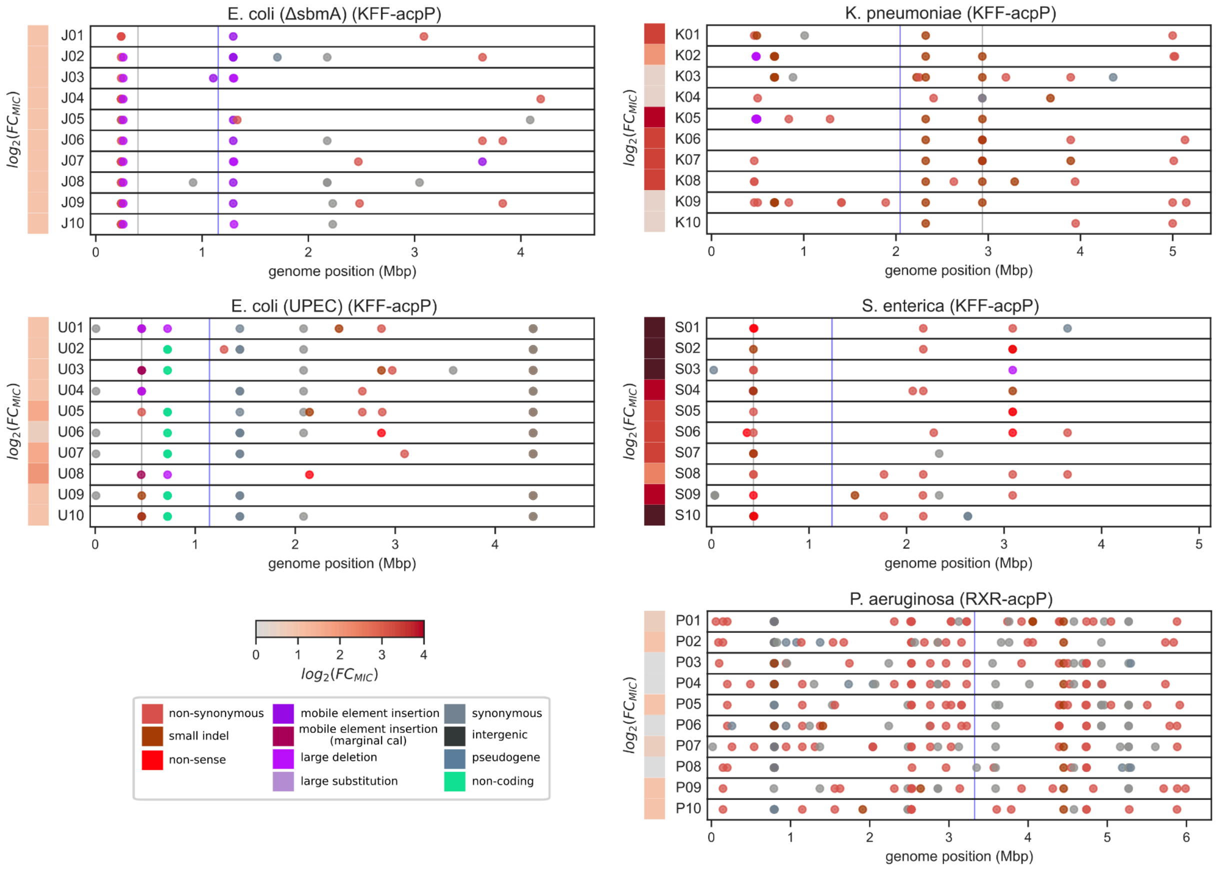


**Supplementary Figure 4. Genome-wide distribution of genetic variants in each biological replicate for those species/ASO formulations not shown in Figure 3.** The MIC fold-change with respect to the ancestral isolate is shown in gray-to-red heatmaps on the left. The gray and blue vertical lines indicate the location of the *sbmA* and the PNA target gene (*acpP*), respectively.

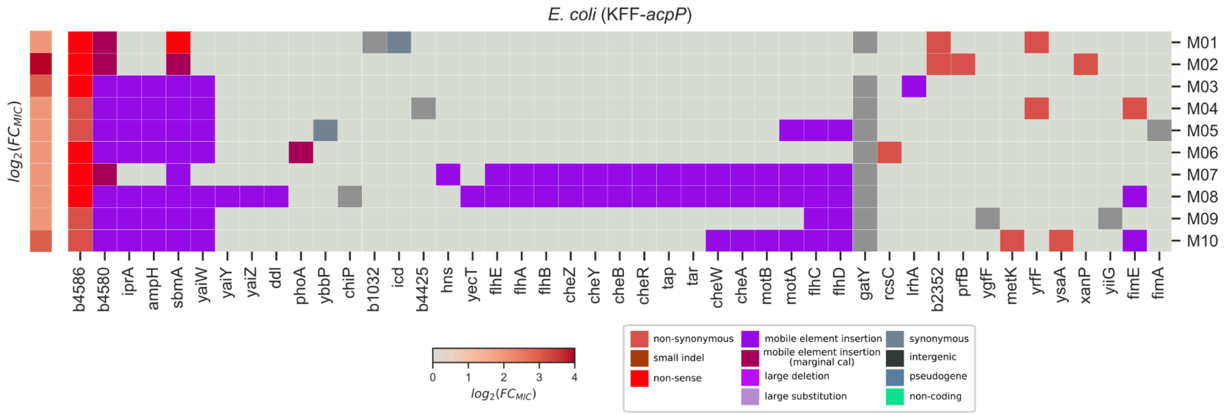


**Supplementary Figure 5. Genome-wide distribution of genetic variants in *E. coli* adapted to KFF-*acpP*.** Genes are ordered according to their position in the chromosome. Light gray boxes indicate that the gene does not have a genetic variant in a sample. The MIC fold-change with respect to the ancestral isolate is shown in gray-to-red heatmaps on the left.

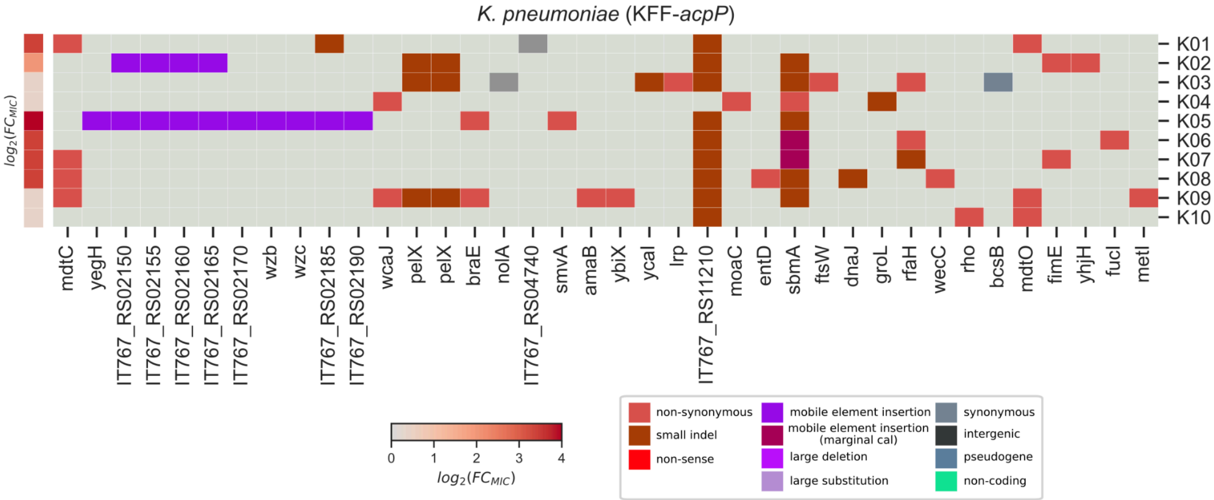


**Supplementary Figure 6. Genome-wide distribution of genetic variants in *K. pneumoniae* adapted to KFF-*acpP*.** Genes are ordered according to their position in the chromosome. Light gray boxes indicate that the gene does not have a genetic variant in a sample. The MIC fold-change with respect to the ancestral isolate is shown in gray-to-red heatmaps on the left.

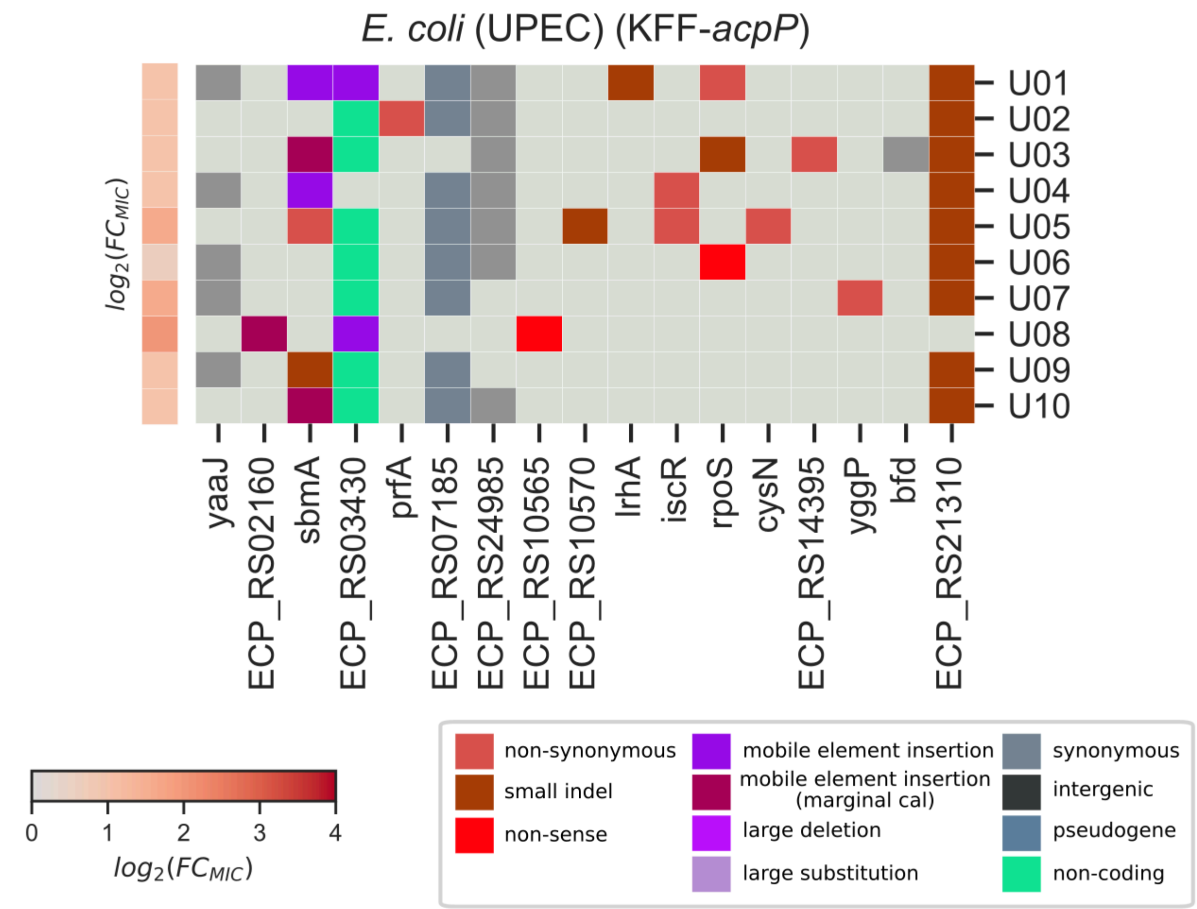


**Supplementary Figure 7. Genome-wide distribution of genetic variants in *E. coli* UPEC adapted to KFF-*acpP*.** Genes are ordered according to their position in the chromosome. Light gray boxes indicate that the gene does not have a genetic variant in a sample. The MIC fold-change with respect to the ancestral isolate is shown in gray-to-red heatmaps on the left.

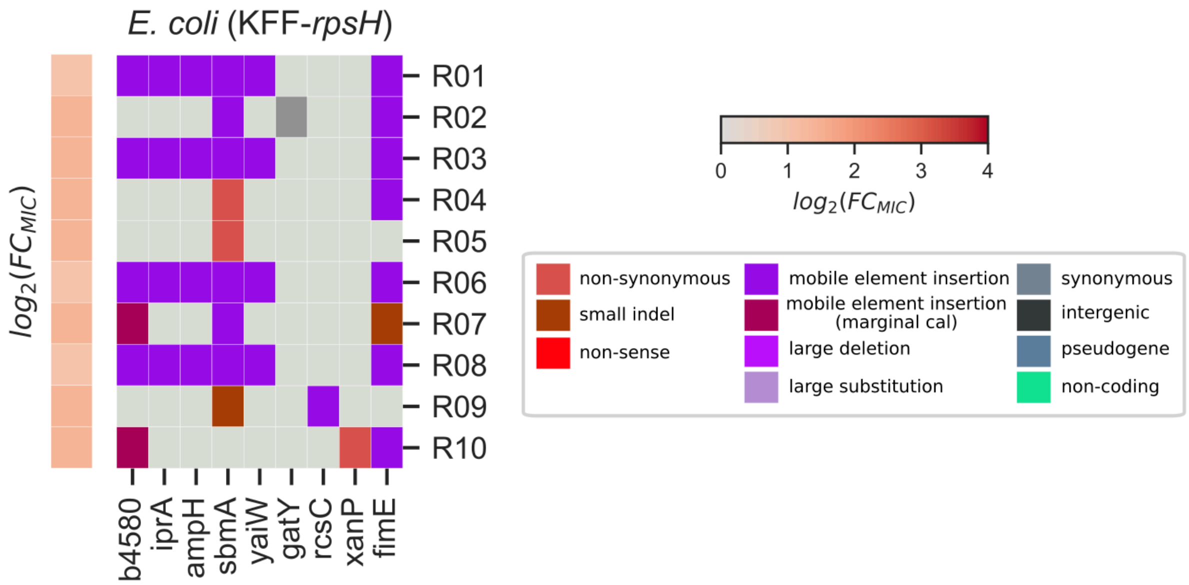


**Supplementary Figure 8. Genome-wide distribution of genetic variants in *E. coli* adapted to KFF-*rpsH*.** Genes are ordered according to their position in the chromosome. Light gray boxes indicate that the gene does not have a genetic variant in a sample. The MIC fold-change with respect to the ancestral isolate is shown in gray-to-red heatmaps on the left.

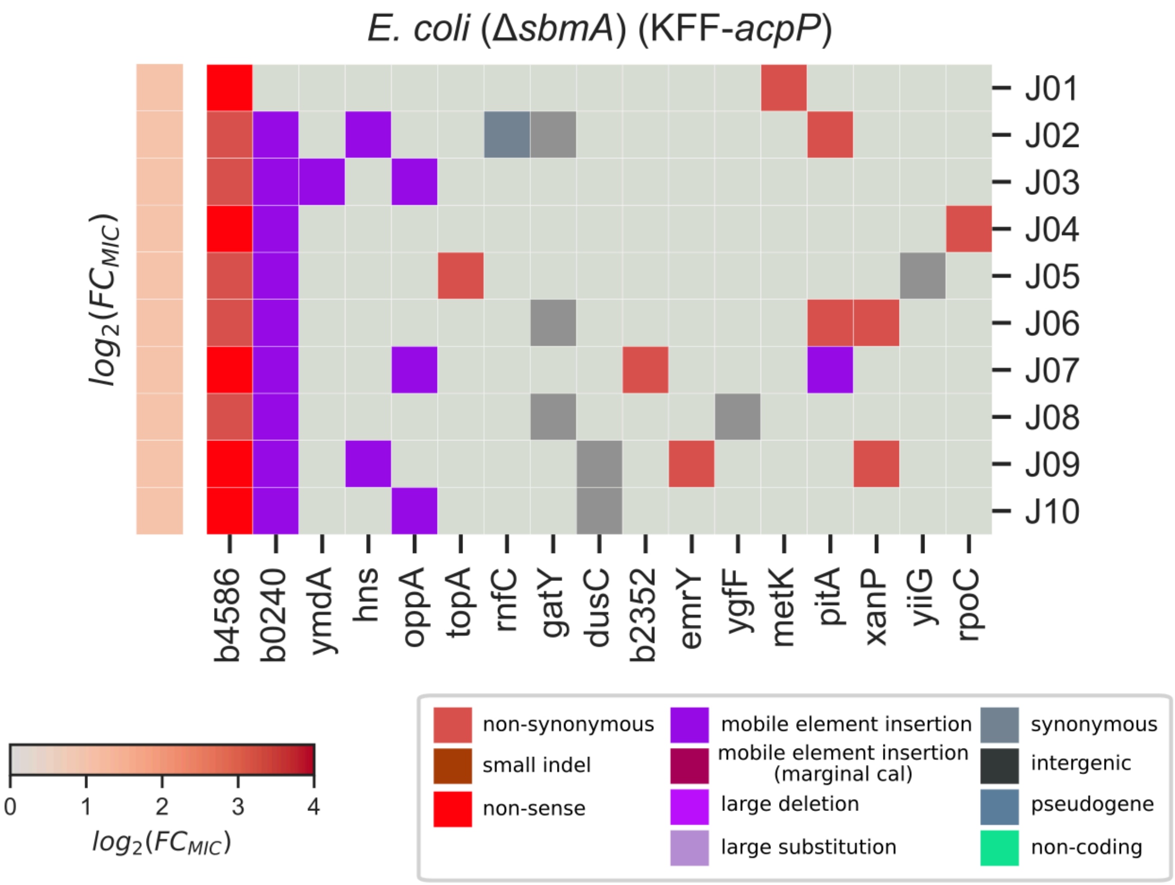


**Supplementary Figure 9. Genome-wide distribution of genetic variants in the *E. coli sbmA* knockout strain adapted to KFF-*acpP*.** Genes are ordered according to their position in the chromosome. Light gray boxes indicate that the gene does not have a genetic variant in a sample. The MIC fold-change with respect to the ancestral isolate is shown in gray-to-red heatmaps on the left.

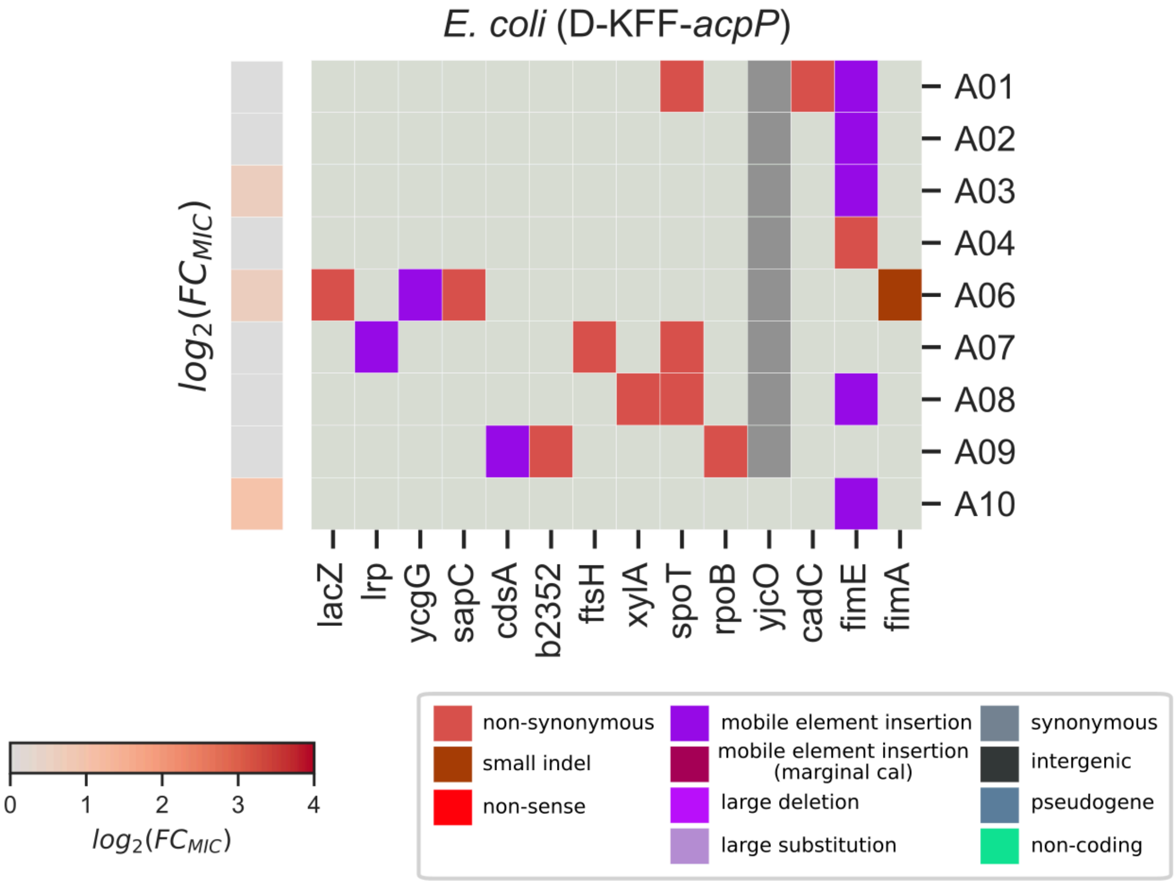


**Supplementary Figure 10. Genome-wide distribution of genetic variants in *E. coli* adapted to D-KFF-*acpP*.** Genes are ordered according to their position in the chromosome. Light gray boxes indicate that the gene does not have a genetic variant in a sample. The MIC fold-change with respect to the ancestral isolate is shown in gray-to-red heatmaps on the left.

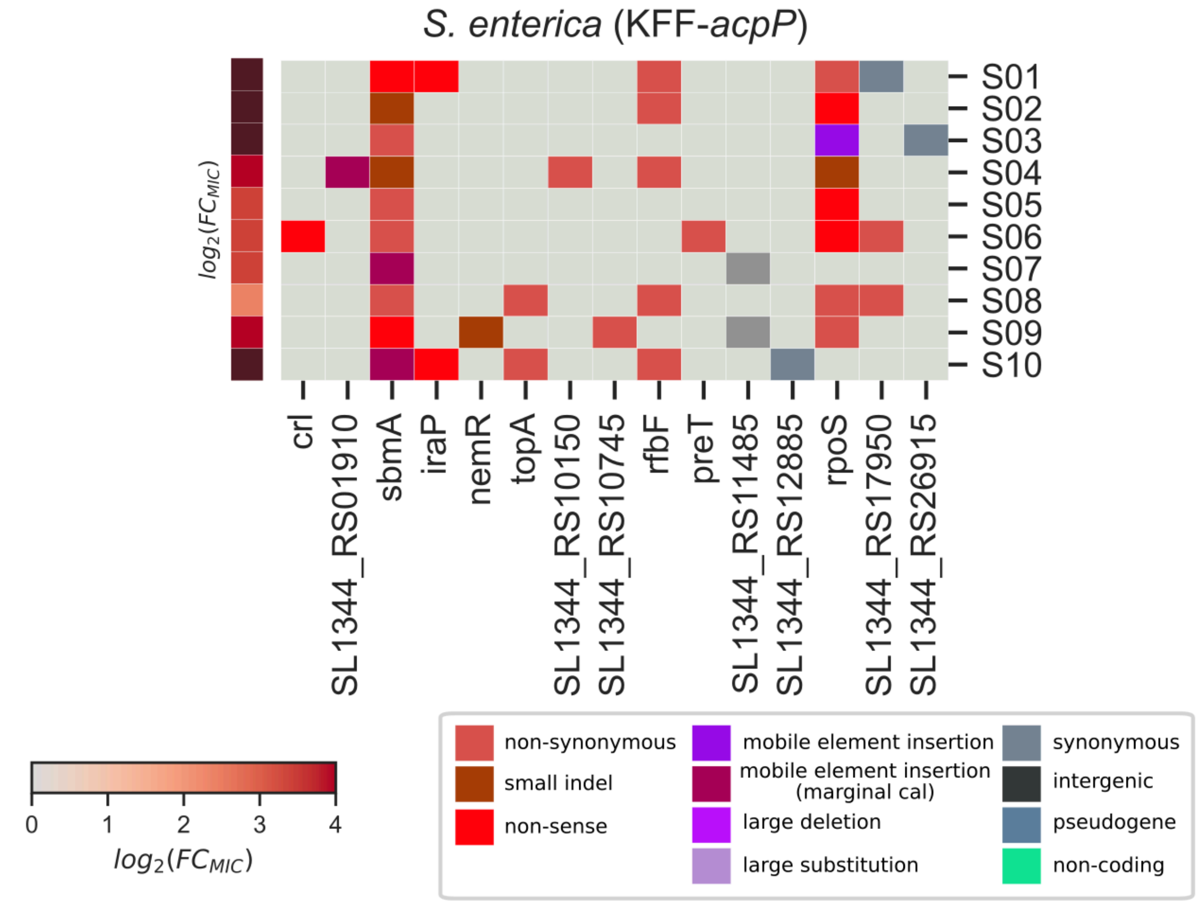


**Supplementary Figure 11. Genome-wide distribution of genetic variants in *S. enterica* adapted to KFF-*acpP*.** Genes are ordered according to their position in the chromosome. Light gray boxes indicate that the gene does not have a genetic variant in a sample. The MIC fold-change with respect to the ancestral isolate is shown in gray-to-red heatmaps on the left.

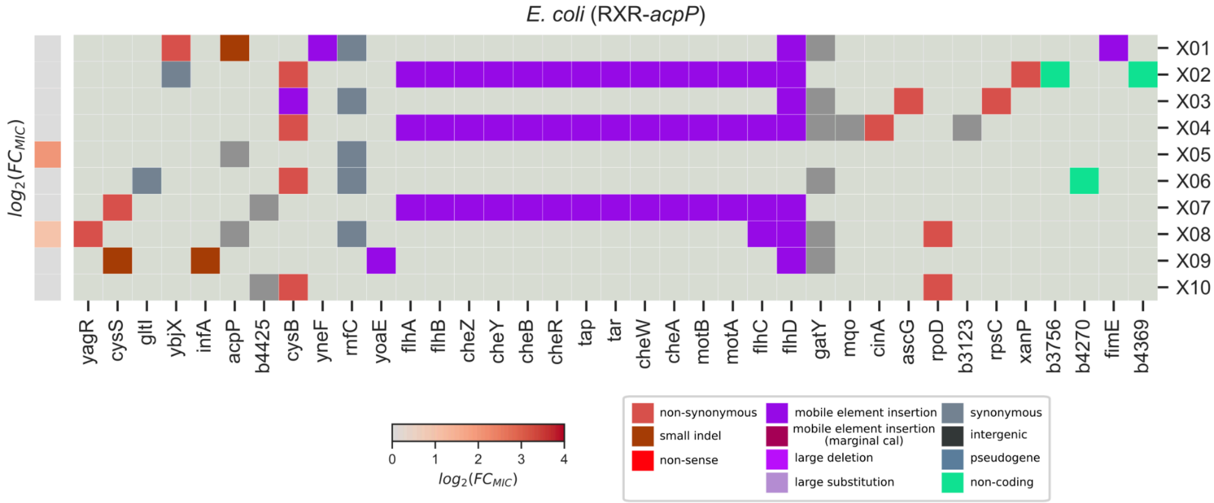


**Supplementary Figure 12. Genome-wide distribution of genetic variants in *E. coli* adapted to RXR-*acpP*.** Genes are ordered according to their position in the chromosome. Light gray boxes indicate that the gene does not have a genetic variant in a sample. The MIC fold-change with respect to the ancestral isolate is shown in gray-to-red heatmaps on the left.

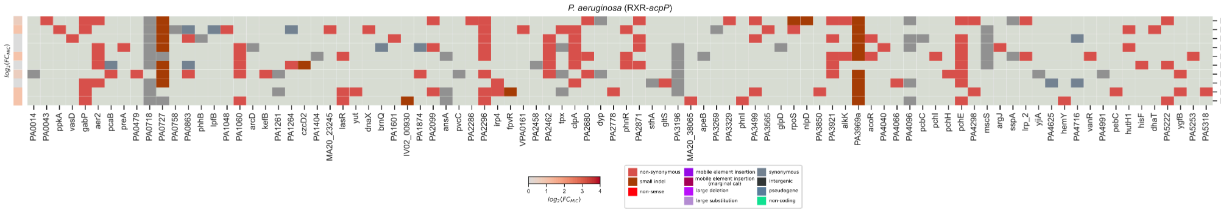


**Supplementary Figure 13. Genome-wide distribution of genetic variants in *P. aeruginosa* adapted to RXR-*acpP*.** Genes are ordered according to their position in the chromosome. Light gray boxes indicate that the gene does not have a genetic variant in a sample. The MIC fold-change with respect to the ancestral isolate is shown in gray-to-red heatmaps on the left.
